# UGDH promotes tumor-initiating cells and a fibroinflammatory tumor microenvironment in ovarian cancer

**DOI:** 10.1101/2022.10.07.509566

**Authors:** Brittney S. Harrington, Rahul Kamdar, Franklin Ning, Soumya Korrapati, Michael W. Caminear, Lidia F. Hernandez, Donna Butcher, Elijah Edmondson, Nadia Traficante, Australian Ovarian Cancer Study Group, Madeline Gough, Rebecca Rogers, Rohan Lourie, Jyoti Shetty, Bao Tran, Fathi Elloumi, Abdalla Abdelmaksoud, Madhu Lal Nag, Krystyna Mazan-Mamczarz, Carrie D. House, John D. Hooper, David D. L. Bowtell, Christina M. Annunziata

## Abstract

Epithelial ovarian cancer (EOC) is a global health burden, with the poorest five-year survival rate of the gynecological malignancies due to diagnosis at advanced stage and high recurrence rate. Recurrence in EOC is driven by the survival of chemoresistant, stem-like tumor-initiating cells (TICs) that are supported by a complex extracellular matrix (ECM) and immunosuppressive microenvironment. To target TICs to prevent recurrence, we identified genes critical for TIC viability from a whole genome siRNA screen. A top hit was the cancer-associated, proteoglycan subunit synthesis enzyme UDP-glucose dehydrogenase (UGDH). Immunohistochemistry was used to delineate UGDH expression in histological and molecular subtypes of EOC. High UGDH expression was observed in the majority of high-grade serous ovarian cancers with variable expression in clear cell, mucinous and endometrioid histotypes. A distinctive prognostic difference was revealed when serous cancers were stratified by molecular subtype, where high UGDH was associated with poor prognosis in the C1/Mesenchymal subtype and low UGDH was associated with poor prognosis in the C4/Differentiated subtype. Ovarian cancer cell lines were subtyped according to the molecular subtypes, and we examined the effect of modulating UGDH expression in cell lines representing the C1/Mesenchymal subtype and C4/Differentiated subtypes. Knockdown of UGDH in the C1/Mesenchymal subtype reduced spheroid viability, sphere-formation and the CD133+/ALDH ^high^ TIC population. Conversely, overexpression of UGDH in the differentiated subtype enhanced spheroid formation but reduced the TIC population. Inflammatory cytokine expression was altered by UGDH expression. In co-culture models, altering UGDH expression in spheroids affected the gene expression of mesothelial cells causing changes to matrix remodeling proteins. The effect of UGDH knockdown or overexpression in the C1/Mesenchymal and C4/Differentiated subtypes, respectively, was tested on mouse intrabursal xenografts and showed dynamic changes to the tumor stroma. Knockdown of UGDH reduced tumor burden in C1/Mesenchymal xenografts compared to controls. These data show that modulation of UGDH expression in tumors influences cells in the microenvironment and reveals distinct roles for UGDH in the mesenchymal and differentiated molecular subtypes of EOC. UGDH is a potential therapeutic target in TICs, for the treatment of metastatic and recurrent EOC, particularly in patients with the mesenchymal molecular subtype.

## Introduction

Epithelial ovarian cancer (EOC) remains the most lethal gynecologic malignancy, with 19,880 new cases and 12,810 deaths estimated in the United States in 2022 (1). EOC is defined by a high level of heterogeneity, diagnosis at an advanced stage, and a high rate of disease relapse (2). Five-year survival rates of Stage 1 disease are as high as 90% (2). However, metastasis, often to the omentum and peritoneum, complicates treatment and dramatically reduces survival rates to 30% (3,4). Stratification of high-grade EOCs by molecular subtype reveals differences in survival, disease burden and surgical complexity. For example, the mesenchymal molecular subtype of ovarian cancer has the worst overall survival and is associated with poorer surgical outcomes due to increased upper abdominal metastases, suboptimal debulking and severe postoperative complications (5–7).

The presence of malignant ascites allows dissemination of EOC tumor cells as spheroids to other peritoneal and abdominal sites (8). EOC spheroids harbor stem-like tumor-initiating cells (TICs) and present significant challenges to successful therapy of metastatic EOC as they promote chemoresistance and disease recurrence (9–11). Furthermore, the complex and immunosuppressive tumor microenvironment (TME) of EOC presents significant challenges to treatment and promotes survival and metastasis of TICs (12). Extracellular matrix (ECM) proteoglycans abundant in the EOC TME promote metastasis, bind to and moderate the activity of cytokines and chemokines, and modulate the interactions between heterotypic cell types (13).

We hypothesize that TICs, supported by this complex TME, are a target for therapeutic eradication. In this study, we identified genes essential for spheroid survival and investigated the enzyme UDP-glucose-6 dehydrogenase (UGDH). Functionally, UGDH promotes the synthesis of glycosaminoglycans and proteoglycans which help maintain the integrity of the extracellular matrix (14,15). UGDH produces the substrates necessary for hyaluronic acid by oxidizing the nucleotide sugar UDP-glucose, to UDP-glucoronate (UDP-GlcUA) (16) and is involved in drug and hormone metabolism through glucuronidation (17,18). UGDH has been associated with promoting cancers of the lung (19,20), glioblastoma (21,22), colon (23), prostate (24), breast (25–27) and ovary (28).

Here we examined the expression and localization of UGDH in tissue microarrays of EOC histotypes mucinous, endometrioid, clear cell and serous, as well as in the molecular subtypes of high-grade cancers (29) and report its prognostic value. We show UGDH promotes TIC survival and that targeting this enzyme in the highly aggressive mesenchymal molecular subtype reduces viability post-chemotherapy *in vitro* and tumor growth *in vivo*. Further, alteration of UGDH in spheroids influenced the gene expression of mesothelial cells in co-culture, remodeling the ECM and TME. UGDH is a potential therapeutic target in TICs for the treatment of metastatic and recurrent EOC, especially of the mesenchymal subtype.

## Results

### Identification of UGDH as a functional target in EOC spheroids

Previously, we studied EOC TICs and defined characteristics that promote survival such as enhanced drug metabolism and oxidative stress management and identified drugs targeting TICs that could prevent relapse *in vitro* and *in vivo* (30). Here, we sought to identify novel targets that functionally regulate EOC TICs and performed a whole-genome siRNA functional screen for targets that preferentially reduced viability of EOC TICs. We used the TIC-enriching spheroid culture conditions that we described previously (31) compared to adherent culture. We chose the OV90 cell line as it is *TP53* mutant, homologous recombination repair proficient, BRCA wild-type and resistant to platinum and PARP inhibitors (32,33).

OV90 adherent cells or spheroids were transfected with at least 2 siRNAs per gene and viability was measured after 96 hours. Using the *Z*-score of the viability of spheroids minus adherent cells, we ranked the genes that reduced spheroid viability compared to adherent, with the top 20 highlighted (Figure 1A). To further refine the candidate genes, we examined their expression using RNAseq, in OV90 adherent and spheroid cells and plotted the p-value and fold change for spheroid compared to adherent values (Figure 1B). Five genes with significant p-values (<0.05) and enhanced or consistent expression in the OV90 cells were investigated for mRNA expression in ovarian cancer using data from The Cancer Genome Atlas (TCGA) (34) (Figure 1C). Interestingly, three of these genes: Glutathione transferase α4 (GSTA4), Nicotinamide phosphoribosyltransferase (NAMPT) and UGDH are enzymes with roles in metabolism and detoxification which we had previously shown to be targetable pathways in TIC spheroids (30). We also examined their protein expression in the Human Protein Atlas ((35), proteinatlas.org), and all three had low or no expression in ovarian stromal cells from normal ovarian tissue (n=3), but UGDH and NAMPT expression was significantly increased in ovarian cystadenocarcinoma tissues of mucinous, endometrioid and serous histotypes (n=12) (Figure 1D). GSTA4 is a member of the Phase II detoxifying enzyme superfamily and is associated with liver cancer progression (36). NAMPT regulates intracellular nicotinamide adenine dinucleotide (NAD) levels and cellular metabolism (37). UGDH oxidizes nucleotide sugars to produce the subunits of hyaluronan, an extracellular matrix signaling molecule that is dysregulated in EOC (38). We chose to pursue UGDH due to its medium-high expression in EOC compared to normal ovarian tissue, it appears in multiple GSEA hallmarks enriched in spheroids (30) (Figure 1E) and its reported roles in promoting cancer progression (19–21,25–28).

**Figure 1:**
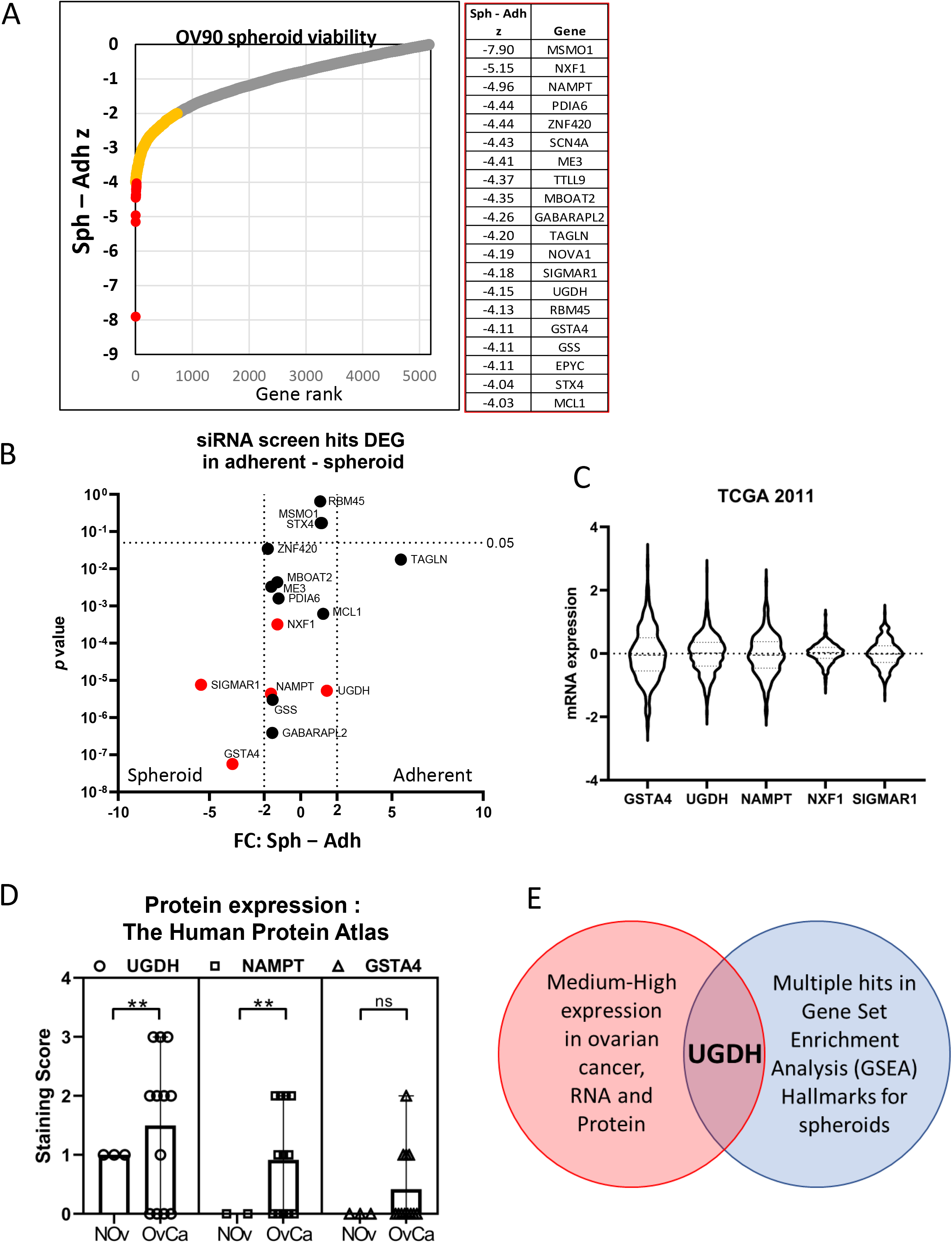
Identification of targets to inhibit the growth and survival of ovarian cancer TICs. A) The top 20 genes identified from an siRNA functional screen that were critical spheroid viability compared to adherent cells using the Z score to compare viability. B) RNA-seq data of OV90 cells cultured as spheroids or cultured adherently from GEO accession number GSE158949. Candidate genes were graphed for gene expression on the x axis and p-value on the y axis. C) mRNA expression of 5 candidate genes in Ovarian Serous Cystadenocarcinoma from TCGA. D) Quantification of protein expression of 3 candidate genes in normal ovarian tissue and ovarian carcinomas from the Huma Protein Atlas. E) Venn diagram to summarize identified target, UGDH.

### UGDH expression in epithelial ovarian cancer histotypes

EOC is a broad description for epithelial malignancies of the ovary and fallopian tube (39). There are 5 main histotypes of EOC: clear cell, mucinous, endometrioid, high-grade serous (HGS) and low-grade serous. These subtypes differ histologically, but also in incidence, disease progression, chemotherapy response and prognosis (39). UGDH was previously detected in mucinous adenocarcinoma and clear cell ovarian cancer tissues by immunohistochemistry (IHC) analysis and was not detected in normal adjacent tissue (28). Here, we sought to characterize UGDH expression in curated tissue microarrays of HGS, endometrioid, mucinous and clear cell EOC histotypes and determine whether UGDH expression was prognostic.

The HGS TMA contained 96 patient tissues, sampled from primary and metastatic sites, and UGDH expression was scored based on previous methods (40,41) as negative, weak, moderate, and strong for both cytoplasmic and nuclear localization (Figure 2A-D). There was a high percentage of positive staining detected overall with only 2.5% of cases being scored as negative for cytoplasmic staining and 11.4% negative for nuclear staining. The distribution of staining intensity and localization in primary and metastatic sites were similar (Figure 2E, F). Correlative analyses were performed on UGDH expression and clinicopathological data, for overall survival and progression-free survival analysis (Supplementary Table 1). We found that localization was not prognostic for HGS (Figure 2G-J). Nuclear expression in HGS was not associated with progression or survival as it was reported for lung adenocarcinoma (19). Although cytoplasmic UGDH expression was associated with survival outcome (Supplementary Table 1), it was not prognostic for overall or progression-free survival (Figure 2G-J). Further investigation with more samples to increase the numbers of HGS cases that were negative for or weakly expressed UGDH may reveal a prognostic effect.

**Figure 2:**
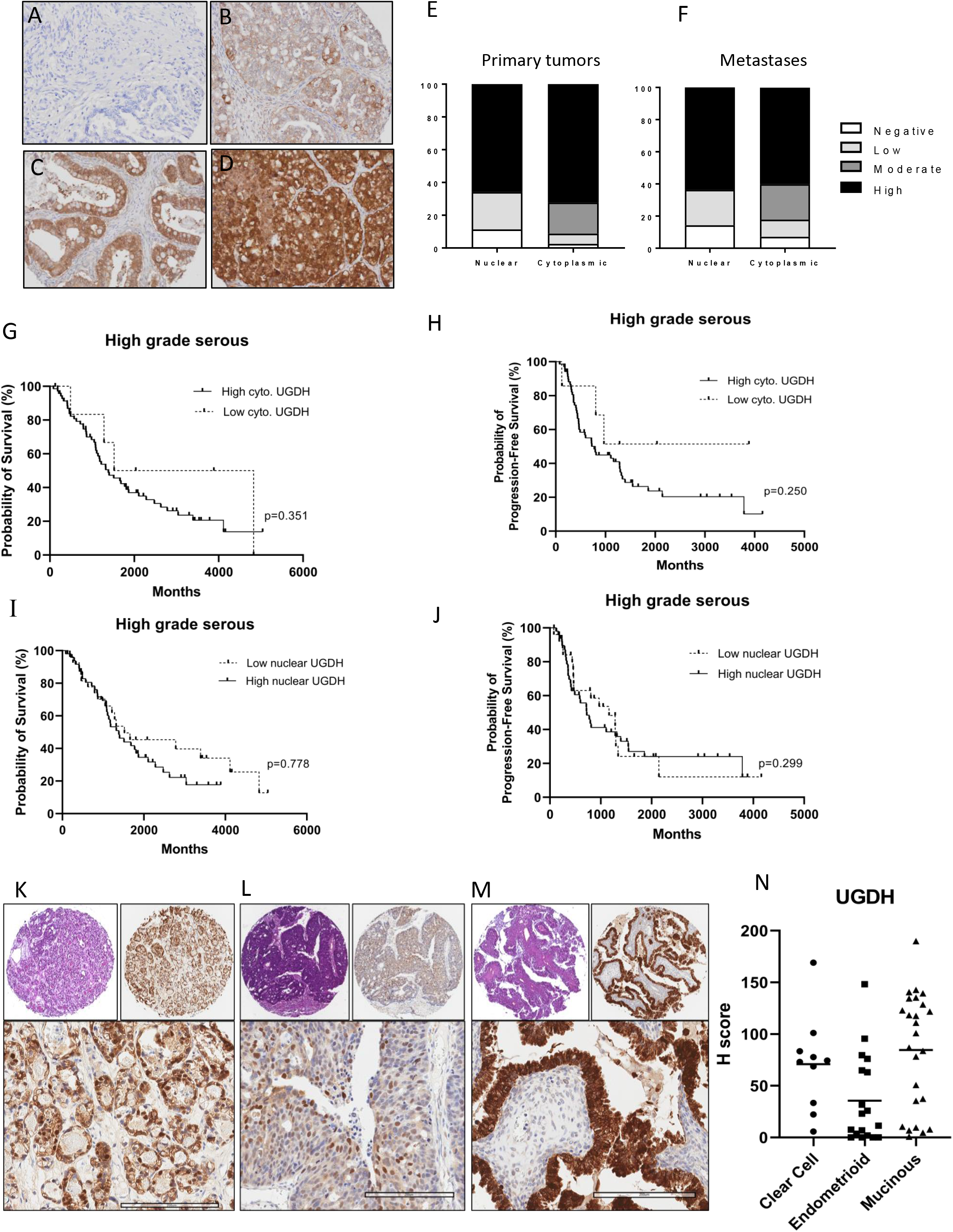
UGDH expression in ovarian cancer histotypes. Representative images of UGDH expression in high grade serous ovarian cancers that were scored as A) Negative, B) Low, C) Moderate or D) Strong, for both cytoplasmic and nuclear localization. E) Proportions of staining scores for cytoplasmic and nuclear expression of UGDH in primary tumors and F) metastases. G) Survival analysis of HGS cancers comparing low versus high cytoplasmic UGDH. H) Survival analysis of HGS cancers comparing low versus high nuclear UGDH. I) Progression-free survival analysis of HGS cancers comparing low versus high cytoplasmic UGDH. J) Progression-free survival analysis of HGS cancers comparing low versus high nuclear UGDH. K) Clear cell, Stage 3C, top left panel H&E in 4X, top right panel IHC in 4X, lower panel IHC 20x, L) Endometrioid, stage 3C, top left panel H&E in 4X, top right panel IHC in 4X, lower panel IHC 20x. M) Mucinous Stage 3C top left panel H&E in 4X, top right panel IHC in 4X, lower panel IHC 20x. N) Expression of UGDH expressed as H-score. Scale bar is 200μm.

For the mucinous, endometrioid and clear cell EOC cases, scoring of UGDH expression was performed using H-score which measures the intensity and proportion of staining (42), (Figure 2K-M). There was abundant UGDH expression in these subtypes, with the highest median expression seen in the mucinous subtype (Figure 2N). UGDH localization did not have prognostic value in these subtypes either. Using the median H score, cases were classified as having higher or lower than the median expression for clinicopathological analyses (43), (Supplementary Table 2). In endometrioid EOC cancers, UGDH expression was associated with stage, where UGDH expression was higher at lower disease stages while expression decreased at higher stage. The number of cases were not sufficient to robustly show differences in overall survival or progression-free survival based on UDGH expression or localization for these subtypes. Of note, most of the cases in the clear cell and mucinous subtypes were International Federation of Gynecology and Obstetrics (FIGO) stage 1 and 2 cancers (44), which typically have a higher survival and lower recurrence rate (45).

### High UGDH expression is associated with poor prognosis in the C1/Mesenchymal molecular subtype

In prior work, Tothill *et. al* profiled the gene expression of 285 serous and endometrioid ovarian, fallopian tube, and peritoneal cancers, as well as a small number of low-grade, low malignant potential tumors. The tumors were categorized into six molecular subtypes (29). The high-grade cancers clustered into subtypes designated C1, C2, C4 and C5, while the C3 and C6 subtypes clustered the low grade, early stage, and low malignant potential tumors (29). The subtypes were characterized by gene expression, histology, immune infiltration, stromal desmoplasia and prognosis, which showed that the C1 and C5 subtypes correlated with a poorer overall and progression-free survival compared to the other subtypes (29). Molecular subtypes for EOC were later characterized independently by the TCGA consortium, and were characterized as Mesenchymal, Immunoreactive, Differentiated and Proliferative (34). Here, we examined UGDH expression in the same TMA used by Tothill *et. al* and describe the subtypes using both the Tothill *et. al* and TCGA designations: C1/Mesenchymal (C1/MES), C2/Immunoreactive (C2/IMR), C4/Differentiated (C4/DIF), C5/Proliferative (C5/PRO).

UGDH expression was highest in the C1/MES subtype, followed by the C5/PRO subtype (Figure 3A, B). The C1/MES subtype has the poorest prognosis of the subtypes (29) and high UGDH expression correlated with shorter overall survival (Figure 3C), but not progression (Supplementary Table 3). The C1/MES subtype was classified by a high stromal signature, with gene expression increases in ECM proteins, proteoglycans and histologically a high level of desmoplasia (29). Interestingly, the C4/DIF subtype that is classified as a low stromal signature but increased immune infiltration, showed the opposite prognostic result for UGDH expression. Low UGDH expression was associated with a significantly poorer overall survival and progression-free survival in this subtype (Figure 3D). The C2/IMR and C5/PRO subtypes did not show significant correlations of UGDH expression with prognosis (Figure 3E, F). We also examined nuclear and cytoplasmic localization of UGDH in the molecular subtypes for prognostic value (Supplementary Table 3). Cytoplasmic and nuclear expression was similar among the cases, in that the cases with high UGDH expression had both high cytoplasmic and nuclear expression and was not prognostic. In the C4/DIF subtype however, the prognostic effect of high UGDH expression tended to be related to cytoplasmic rather than nuclear expression (Supplementary Table 3).

**Figure 3:**
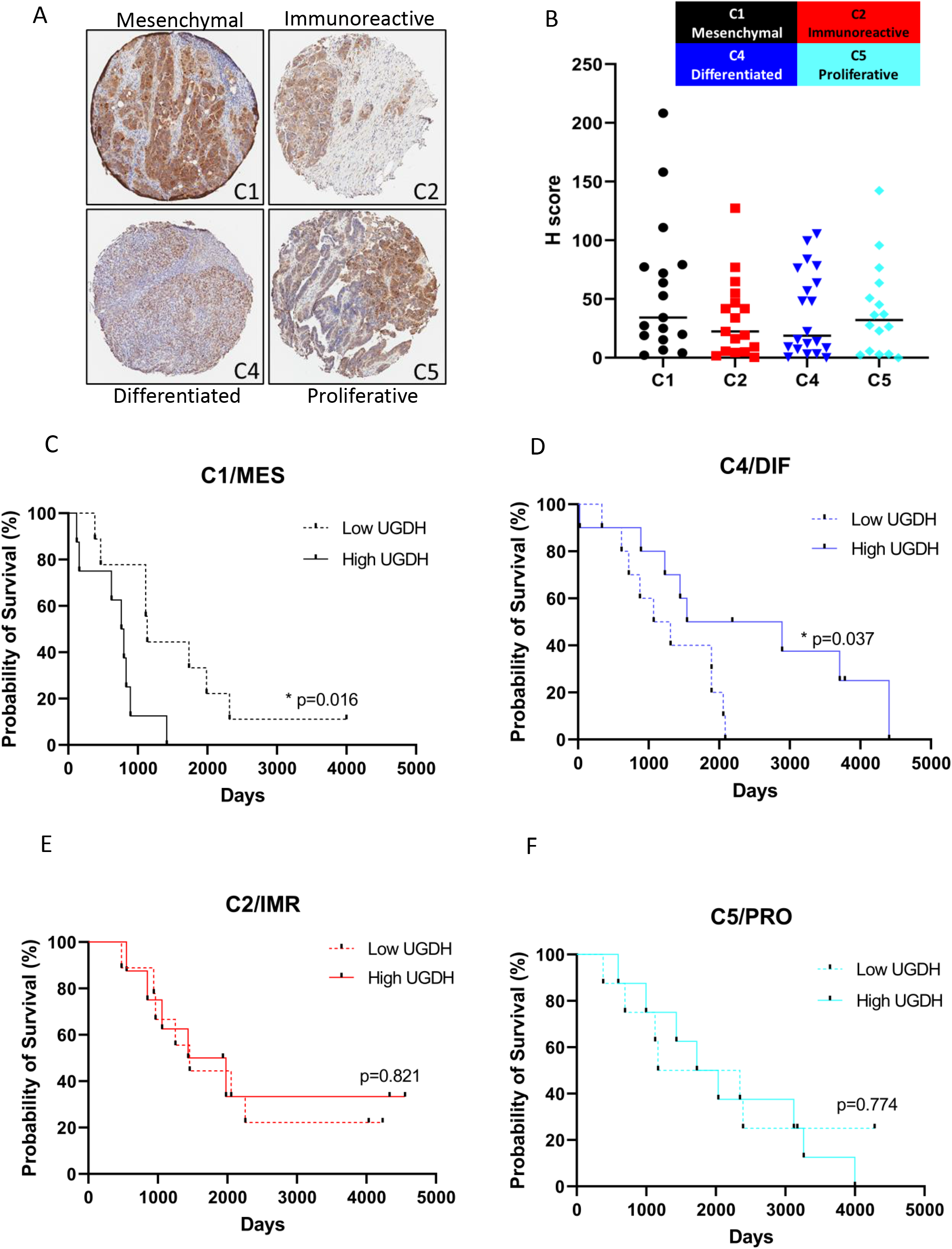
UGDH expression in molecular subtypes of high grade epithelial ovarian cancers. A) Representative images of UGDH expression in TMA cores from molecular subtypes C1, C2, C3 and C4 at 4X magnification. B) Expression of UGDH expressed as H-score. C) Survival analysis of C1 subtype comparing low versus high UGDH H-score (above or below the median). D) Survival analysis of C4 subtype comparing low versus high UGDH H-score (above or below the median). E) Survival analysis of C2 subtype comparing low versus high UGDH H-score (above or below the median). F) Survival analysis of C5 subtype comparing low versus high UGDH H-score (above or below the median).

### UGDH expression in cell lines clustered by molecular subtyping analysis

To model the molecular subtypes *in vitro* we classified ovarian cancer cell lines into the molecular subtypes originally annotated by two independent datasets of primary cancer specimens (29,34). Molecular subtyping of ovarian cancer cell lines was previously reported, using a different clustering method that classified novel molecular subtypes (46). However, we sought to identify cell lines to represent the subgroups in which we identified prognostic implications for UGDH. Therefore, we clustered the cell lines using four clusters representing the four molecular subtypes C1/MES, C2/IMR, C4/DIF, and C5/PRO with biological duplicates (Figure 4A). Two representative cell lines for each subtype were examined for expression of UGDH from both adherent and TIC spheroid culture conditions by Western blot analysis (Figure 4B). UGDH expression was highest in OV90 (C1/MES) in both culture conditions, and notably OVCAR3 in the C5/PRO subtype showed elevated expression in the TIC spheroid culture condition. UGDH expression in the cell lines did resemble the finding of the IHC performed on patient samples of the molecular subtypes, where the C1 subtype tumors had the highest median H-score for UGDH, followed by the C5/PRO subtype and lower expression in the C2/IMR and C4/DIF subtypes. From this analysis, we used OV90 to represent the C1/MES subtype, in which high UGDH expression correlated with poorer survival, and ACI23 to represent the C4/DIF subtype, in which low UGDH expression correlated with shorter survival.

**Figure 4:**
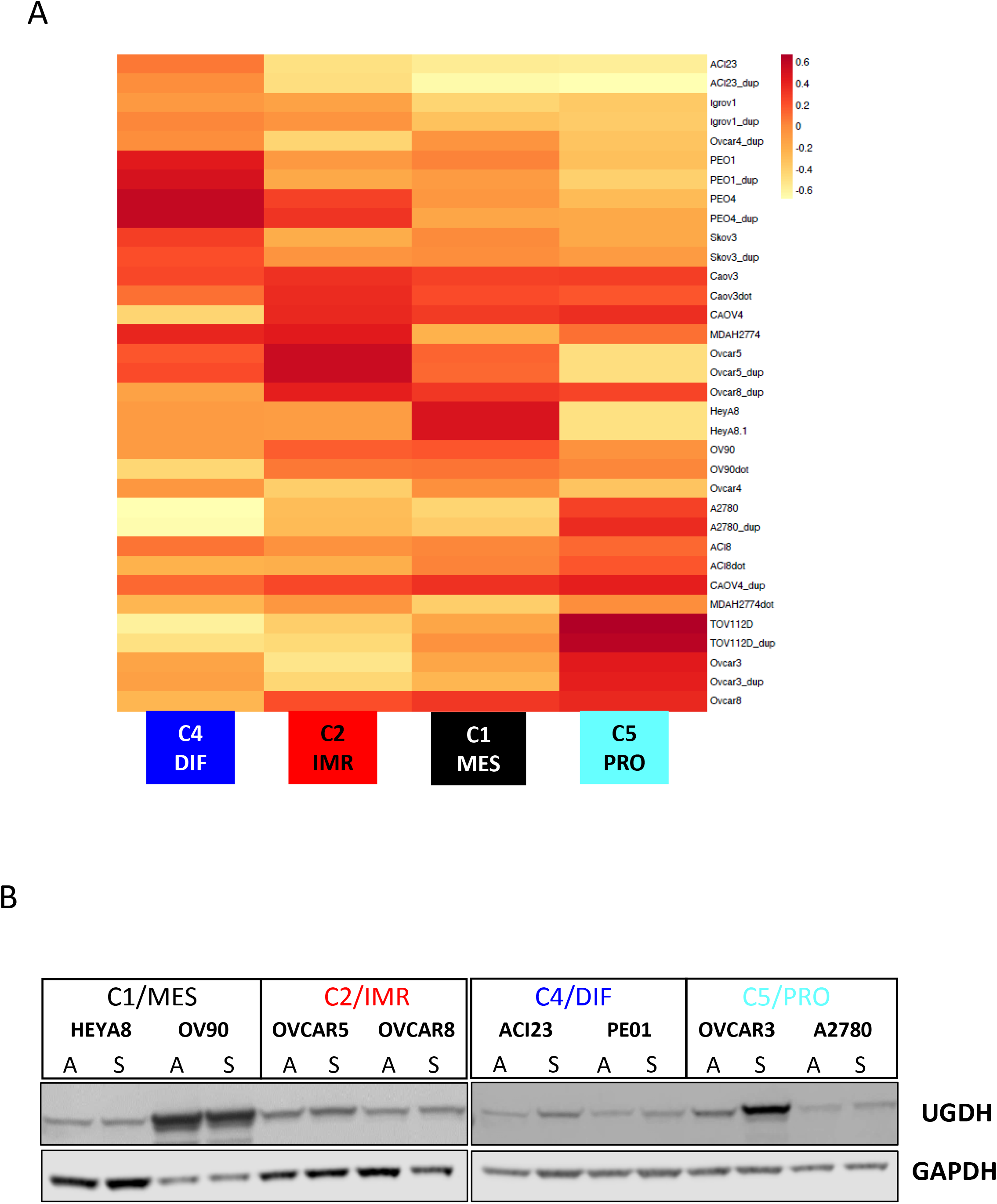
Ovarian cancer cell lines clustered into molecular subtypes examined for UGDH expression. A) Heatmap of cell lines aligned with molecular subtypes. B) Expression of UGDH in cell lines in adherent and spheroid culture conditions by Western blot analysis.

### Spheroid viability is affected by UGDH expression in C1/MES and C4/DIF cell lines

In the C1/MES molecular subtype, high UGDH expression was associated with poorer prognosis, but the opposite was observed for the C4/DIF subtype. Therefore, in comparing the effect of UGDH expression in these subtypes, we silenced UGDH expression in OV90 cells using inducible shRNA (sh459, sh939) and over-expressed UGDH in ACI23 cells. Western blot analysis and densitometry showed efficient silencing was induced in OV90 after 72 hours of doxycycline treatment (Figure 5A), and over-expression in ACI23 (Figure 5B) in both adherent and spheroid culture conditions compared to controls. UGDH knockdown in OV90 changed the morphology of adherent cultures to appear more epithelial and ‘ cobblestone’ like (Figure 5C); in contrast, overexpression of UGDH in ACI23 made the cells less differentiated and more mesenchymal in appearance (Figure 5D). The viability of cells in adherent and spheroid culture conditions was examined to validate the findings from the initial siRNA screen of OV90 cells, identifying UGDH as a potential target. In adherent conditions, alteration of UGDH expression did not significantly affect viability of either cell line, but induction of knockdown in formed OV90 spheroids significantly reduced viability by 48-60% (Figure 5E), confirming the effect observed in the siRNA screen. Overexpression of UGDH increased spheroid viability of ACI23, compared to vector control (Figure 5F). The effect of modulating UGDH expression was also examined on spheroid formation, where UGDH silencing was induced from the time of plating. In OV90, UGDH silencing significantly reduced spheroid formation compared to the negative control (Figure 5G). Over-expression of UGDH in ACI23 however, increased the number of spheres compared to the vector control (Figure 5H). These data show that on spheroid viability and formation are greatly reduced when UGDH is silenced in the C1/MES subtype and while these phenotypes are enhanced when UGDH is overexpressed in the C4/DIF subtype.

**Figure 5:**
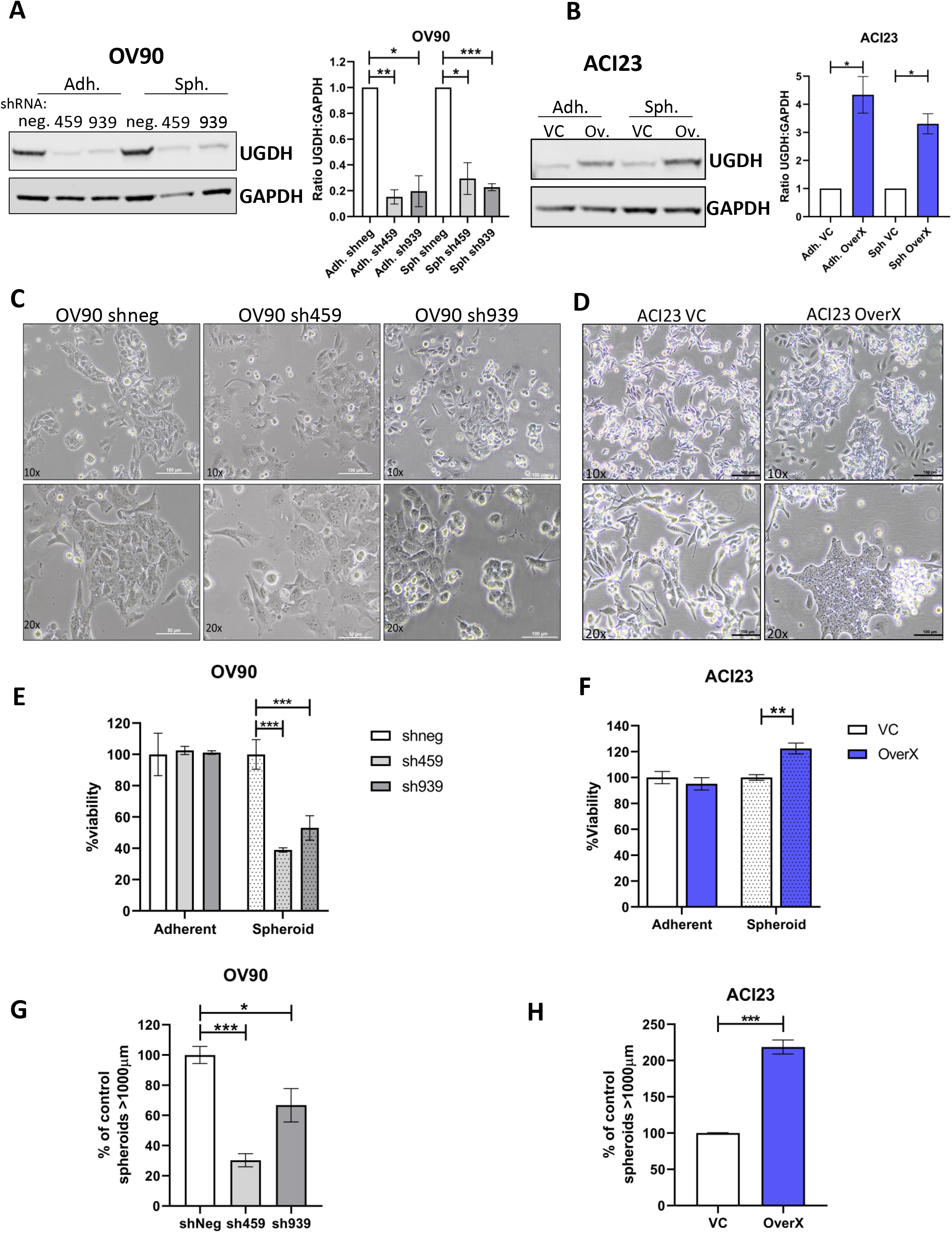
Effects of UGDH knockdown in OV90, and over-expression in ACI23 *in vitro*. Western blot and densitometry analysis of UGDH expression in A) OV90 cells with doxycycline inducible negative control shRNA (shneg) or doxycycline inducible shRNA targeting UGDH (sh459, sh939) after 3 days of doxycycline induction in indicated culture conditions and quantified by densitometry B) ACI23 cells with stably expressed vector control (VC) or UGDH (Ov, OverX) grown for 3 days in indicated culture conditions and quantified by densitometry. C) OV90 shneg, sh459 and sh939 and D) ACI23 VC and OverX cells representative brightfield images of adherent cell culture morphology. Cell viability of E) OV90 shneg, sh459 and sh939 and F) ACI23 VC and OverX grown in adherent or spheroid conditions. Sphere formation capacity of G) OV90 shneg, sh459 and sh939 and H) ACI23 VC and OverX cells. *p<0.05, **p<0.01, ***p<0.001. Scale bar is 100μm.

### UGDH silencing in C1/MES, and over-expression in C4/DIF, reduces TICs in vitro

The spheroid culture condition enriches for the TIC population in ovarian cancer cell lines, which cause enhanced tumor growth in mouse models and promote relapse (30,31,47). Therefore, we examined whether targeting UGDH could affect the features of TICs including colony formation, expression of stem cell markers, and relapse *in vitro*. The colony forming capacity of OV90 cells was significantly reduced by UGDH knockdown compared to the negative control, but over-expression in ACI23 caused no significant difference (Figure 6A, 6B). We and others have shown that the CD133+/ALDH ^high^ cell population are TICs (31,47,48). Examining these markers in spheroid cultures of OV90 and ACI23 cells with altered UGDH expression revealed that silencing in OV90 cells (Figure 6C), and overexpression in ACI23 cells (Figure 6D) caused a significant reduction in this population compared to controls. The same effect caused by opposing expression of UGDH in the cell lines may be explained by different mechanisms. In OV90 cells, the reduction of viability caused by UGDH knockdown in spheroids may explain the overall reduction in CD133+/ALDH^high^ cells. And in ACI23, overexpression of UGDH may out-compete ALDH for NAD+ substrate, as both are dependent on this for activity (16,49), thus causing reduced ALDH activity to be observed. Finally, we used our previously reported *in vitro* relapse model (30,50) to directly assess the potential for spheroids with altered UGDH to promote growth and persist after chemotherapy. The cell lines were grown adherently for 48 hours and treated with a sub-lethal dose of carboplatin or vehicle; the viable populations remaining after treatment were then cultured in TIC-enriching spheroid conditions and assessed for cell death. Knockdown of UGDH in OV90 spheroids after carboplatin treatment significantly increased cell death, compared to the negative control (Figure 6E). Significantly increased cell death was also observed in ACI23 spheroids generated after carboplatin treatment overexpressing UGDH compared to the vehicle control (Figure 6D). These data indicate that differential UGDH expression is important for the composition of the spheroids and the TIC population that drives recurrence, with opposite effects in C1/MES and C4/DIF subtypes.

**Figure 6:**
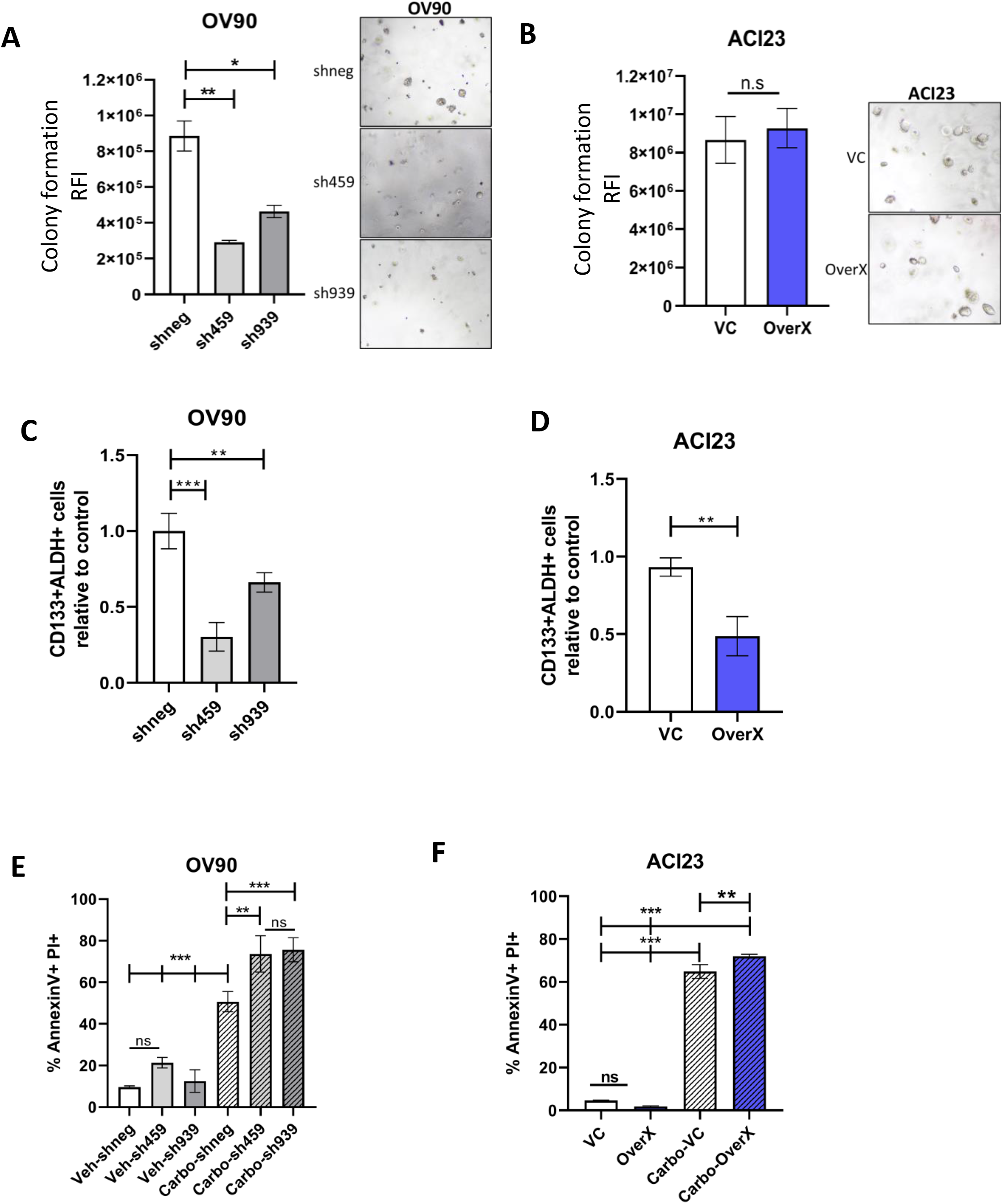
Effects of UGDH knockdown in OV90, and over-expression in ACI23 on TIC populations *in vitro*. A) Colony forming capacity of UGDH knockdown (sh459, sh939) compared to control (shneg) in OV90 cells. B) Colony-forming capacity in ACI23 VC or OverX cells. C) Quantification of the proportion of CD133+ ALDH+ cells in UGDH knockdown (sh459, sh939) compared to shneg OV90 cells grown in TIC-enriching spheroid conditions. D) Quantification of the proportion of CD133+ ALDH+ cells in ACI23 OverX compared to ACI23 VC cells grown in TIC-enriching spheroid conditions. Analysis of cell death by AnnexinV and PI double positive cells from *in vitro* relapse model in spheroids generated from viable cells collected after 48 hours of carboplatin treatment followed by E) induction of UGDH silencing (sh459, sh939) compared to shneg control in OV90 and F) over-expression of UGDH compared to vector control in ACI23. *p<0.05, **p<0.01, ***p<0.001.

### UGDH alters cytokine secretion in spheroids, and gene expression of mesothelial cells in co-culture

In comparing the C1/MES and C4/DIF molecular subtypes, stromal response was the major histological difference between these groups. Therefore, we next examined an *in vitro* model of the peritoneal stroma of EOC by co-culturing mesothelial cell line LP3 with EOC spheroids with altered UGDH expression. We assessed gene expression in UGDH-altered spheroids alone, or in co-culture with UGDH-altered spheroids by qRT-PCR and compared it to mesothelial cells alone (Table 1). When UGDH was knocked down in the OV90 spheroids representing the C1/MES subtype, there was a decrease in the expression of ECM components VCAN and TNC, and increased expression of metalloprotease inhibitor TIMP3, and cell-matrix interacting proteins FN1 and CDH1 (Figure 7A). When these spheroids were co-cultured with LP3, there was a further decrease in VCAN expression, as well as a decrease in matrix remodeling enzyme MMP1 and ECM interacting protein LAMA3 expression. These changes suggest that UGDH knockdown on the C1/MES spheroids causes a decrease in extracellular matrix remodeling and invasive potential due to decreased matrix protease and ECM component expression. We also examined the expression of the same markers in co-cultures of the ACI23 spheroids representing the C4/DIF subtype, with overexpression of UGDH. The overexpression in this subtype replicated some of the effects of knockdown in the C1/MES spheroids, where MMP1 expression was decreased, and TIMP3 expression was increased when UGDH was overexpressed in the spheroids and when in co-culture with LP3 (Figure 7B). However, VCAN expression increased in the overexpressing spheroids and in co-culture. Other changes in this subtype with overexpressed UGDH included reduced COL1A1, FN1 and TGFB expression in co-cultures as well as reduced CDH1 in spheroids. This suggests that the C4/DIF subtype may become more desmoplastic and less differentiated when UGDH is over-expressed. As cytokines can be modulated by the ECM and influence the TME, we were also interested in whether UGDH expression influenced cytokine secretion in the spheroids. In OV90, when UGDH was knocked down, IL-6 and IL-8, levels increased significantly (Figure 7C, D). In the ACI23 cells, when UGDH was overexpressed, there was a significant increase in IL-6, IL-8, and MCP-1 compared to controls (Figure 7E, F, G). These data suggest UGDH differentially influences the tumor microenvironment and regulates inflammatory cytokines in a subtype-specific manner.

**Figure 7:**
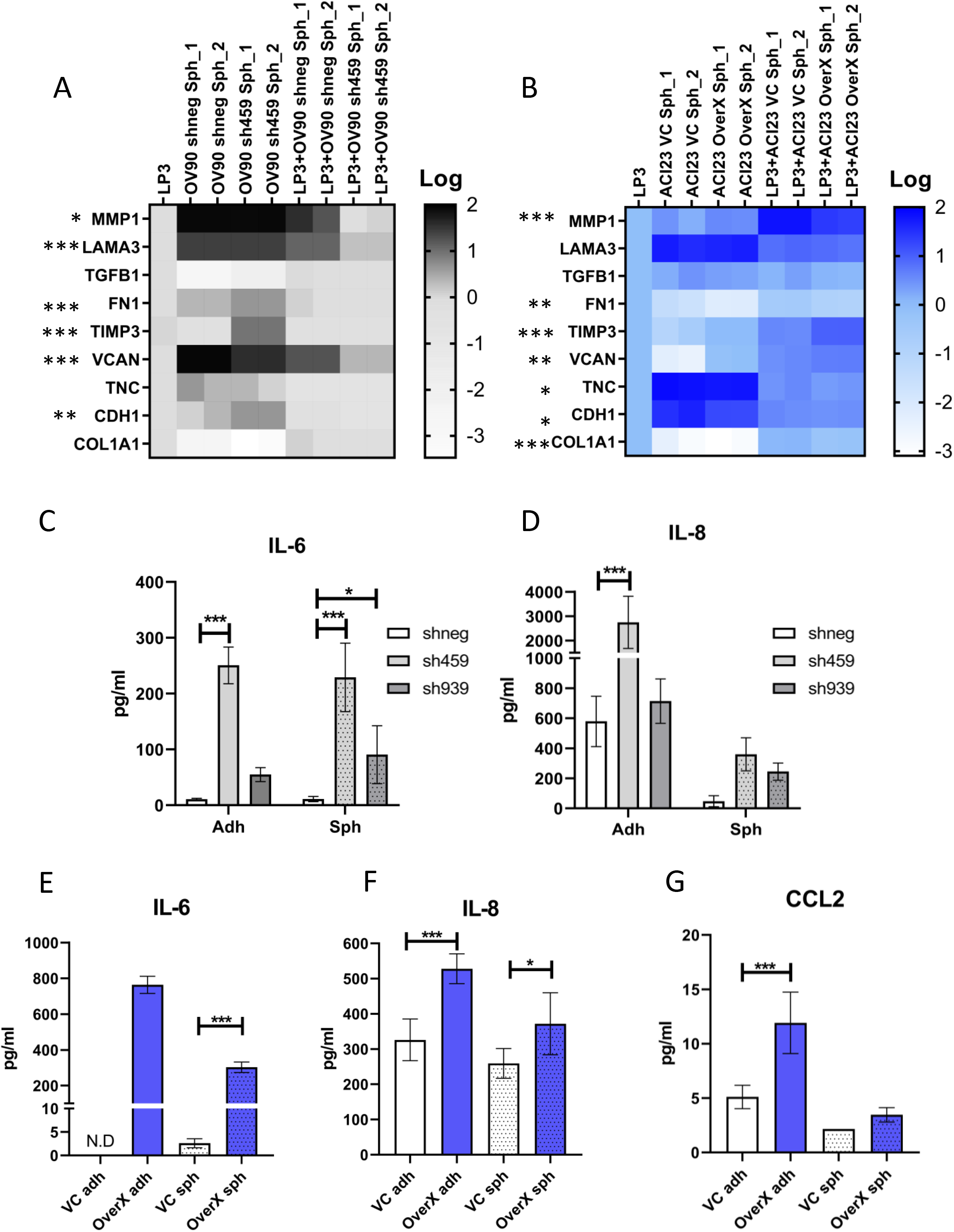
UGDH expression in spheroids alters gene expression of mesothelial cells in co-culture and cytokine expression. Spheroids were generated and knockdown induced with doxycycline before co-culture with LP3 mesothelial adherent monolayers for 24 hours. A) Heatmap of expression of genes altered in OV90 spheroids when UGDH was knocked down and in co-cultures with LP3, relative to LP3 alone. Asterisks indicate statistical significance, given in Table 1. B) Heatmap of expression of genes altered in ACI23 spheroids when UGDH was overexpressed and in co-cultures with LP3, relative to LP3 alone. Asterisks indicate statistical significance, given in Table 1. Expression of C) IL-6 D) IL-8 in supernatant from OV90 adherent cells or spheroids with UGDH knockdown compared to controls. Expression of E) IL-6 F) IL-8 G) MCP-1 in supernatant from ACI23 adherent cells or spheroids with UGDH overexpression compared to controls. *p<0.05, **p<0.01, ***p<0.001.

**Table 1:** UGDH expression in spheroids alters gene expression of mesothelial cells in co-culture. Relative expression of indicated genes compared to LP3 cells alone were compared by one-way ANOVA with Tukey’s multiple comparisons post-test, p values are given. *p<0.05, **p<0.01, ***p<0.001.

|  | **MMP1** | **LAMA3** | **TGFB1** | **FN1** | **TIMP3** | **VCAN** | **TNC** | **CDH1** | **COL1A1** |
| --- | --- | --- | --- | --- | --- | --- | --- | --- | --- |
| OV90 shneg Sph vs sh459 Sph | >0.999 | 0.997 | >0.999 | ***<0.001 | ***<0.001 | ***<0.001 | 0.358 | **0.002 | >0.999 |
| LP3+OV90 shneg Sph vs LP3+OV90 sh459 Sph | *0.036 | ***<0.001 | 0.566 | 0.578 | >0.999 | ***<0.001 | >0.999 | 0.993 | 0.574 |
|  | **MMP1** | **LAMA3** | **TGFB1** | **FN1** | **TIMP3** | **VCAN** | **TNC** | **CDH1** | **COL1A1** |
| ACI23 VC Sph vs ACI23 OverX Sph | 0.833 | 0.997 | 0.997 | 0.183 | **0.002 | *0.047 | *0.041 | *0.021 | >0.999 |
| LP3+ACI23 VC Sph vs LP3+ACI23 OverX Sph | ***<0.001 | 0.999 | 0.861 | **0.001 | ***<0.001 | **0.005 | 0.999 | >0.999 | ***<0.001 |

### UGDH expression in tumor xenografts promotes fibroinflammatory changes in the stroma

The effect of UGDH knockdown in C1/MES and overexpression in C4/DIF was tested on mouse intrabursal xenografts of OV90 and ACI23 cells, respectively. The mice were followed for overall survival to determine if the same prognostic outcome that was observed in the patients could be replicated. In the C1/MES groups, knockdown of UGDH in OV90 xenografts showed a strong trend towards improved survival compared to the negative control OV90 xenografts (Figure 8A). These results replicate the prognostic results of UGDH expression in patients with EOC in the C1/MES molecular subtype. Conversely, overexpression of UGDH in the C4/DIF ACI23 xenografts did not significantly affect survival compared to controls (Figure 8B). The changes to gene expression of co-cultured cells *in vitro* also prompted investigation of the histomorphology of OV90 and ACI23 xenografts. The xenografts of ACI23 and OV90 differed greatly, with ACI23 xenografts manifesting as large, differentiated neoplasms with areas of necrosis within the ovarian bursa. OV90 xenografts showed multiple foci of smaller neoplastic masses in the bursa and some intratumoral hemorrhage (Figure 8 C, D). Within the OV90 xenografts, UGDH knockdown significantly reduced tumor burden compared to controls (Figure 8C). In comparison, overexpression of UGDH in the ACI23 xenografts did not significantly affect tumor size (Figure 8D). The histomorphology of the xenografts was examined for fibrosis and collagen deposition using Masson’s trichrome stain (Figure 8E, F). The small numbers of viable tumor from OV90 xenografts with UGDH knockdown prevented thorough assessment of effects *in vivo*. Tumors with UGDH overexpression showed enhanced collagen deposition but fibrotic stroma (Figure 8F) and increased expression of VCAN, LAMA3 and IL-6 (Figure 8G-I), consistent with *in vitro* co-culture findings. These data indicate that the alteration of UGDH in tumor cells influences the TME to become pro-inflammatory.

**Figure 8:**
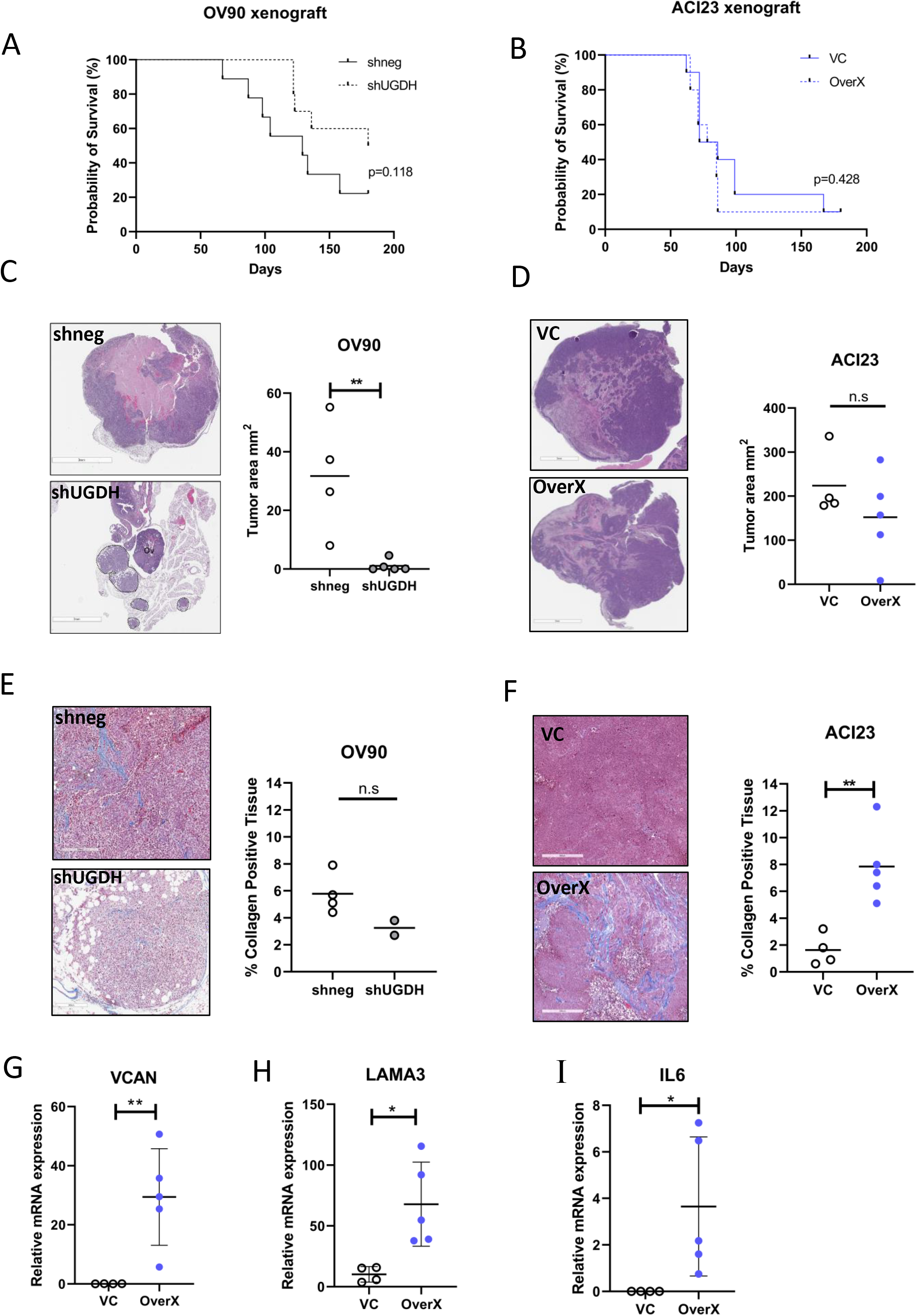
Overall survival, tumor size and histomorphology of xenografts of OV90 with UGDH knockdown, and ACI23 with UGDH over-expression. A) Survival analysis of OV90 xenografts B) Survival analysis of ACI23 xenografts C) H&E images of OV90 xenografts (left) and quantification of tumor size (right). Tumor is marked by dashed lines; ovary is marked Ov. Scale bar is 3mm. D) H&E images of ACI23 xenografts (left) and quantification of tumor size (right). Scale bar is 3mm. E) Massons trichrome staining images of OV90 xenografts (left) and quantification of collagen in the tissue (right) Scale bar is 300um. F) Massons trichrome staining images of ACI23 xenografts (left) and quantification of collagen in the tissue (right). Scale bar is 300um. G) Expression of VCAN, H) LAMA3 and I) IL6 in ACI23 VC and OverX xenograft tumors. *p<0.05, **p<0.01.

## Discussion

The TME of EOC is a complex, immunosuppressive network of heterotypic cell types supported by ECM, cytokines and growth factors and presents a significant challenge to treatment, especially in the mesenchymal molecular subtype. Disease progression and recurrence in EOC is promoted by the TME and the survival of TICs in spheroids, which are targets for therapeutic eradication. We utilized a functional siRNA screen to identify genes essential to spheroid survival and report our findings of UGDH in EOC. UGDH has tumor-promoting functions in multiple cancer types and here we sought to characterize UGDH expression in EOC and identify its roles in supporting TICs and its influence on the TME. We identified key subtype-specific differences indicating that UGDH pro-tumorigenic activity predominates in the mesenchymal subtype of HGS ovarian cancer. This has important implications for the development of therapeutic strategies in this disease.

Previously it was demonstrated that UGDH promoted migration, tumor growth in a subcutaneous model, cell cycle progression and epithelial-mesenchymal transition in ovarian cancer cell lines (28). UGDH expression in mucinous and clear cell EOC subtypes was also examined and found it was elevated compared to normal adjacent tissue. In our extensive range of EOC TMAs we also found strong expression of UGDH in high-grade serous cancers, to such a degree that it was not feasible to correlate with prognoses due to the few cases of negative staining observed. In the clear cell, endometrioid and mucinous tissues we saw a variation in expression, but this was also not indicative of prognoses in the low numbers of cases examined. More cases may provide insight into UGDH as a prognostic marker in these histotypes. We also examined the localization of UGDH in the TMAs to determine if it had a prognostic indication for EOC, similar to what was reported for lung adenocarcinoma (19). In lung adenocarcinoma positive nuclear UGDH localization correlated with lymphatic and vascular invasion, larger tumor size, higher stage, and poor differentiation (19). However, we did not find any correlation between clinicopathological data and UGDH localization in our samples where most samples were positive for both nuclear and cytoplasmic localization. This suggests that in EOC, the function of UGDH in promoting cancer progression is not linked to distinct nuclear or cytoplasmic roles. Moreover, the most significant prognostic indication of UGDH expression was found in the molecular subtypes of EOC. We showed that UGDH expression correlated with prognosis in the molecular subtypes C1/MES and C4/DIF which have distinct stromal phenotypes in terms of histology and immune infiltration (29). Importantly, high UGDH expression had opposite effects in these subtypes. This finding suggests that if therapies were designed to block UGDH activity, they should be specifically directed to women with the mesenchymal subtype and not the differentiated type of HGS.

The C1/MES molecular subtype was described as high stromal reactive, with extensive desmoplasia and immune infiltration within the stroma but lower intertumoral infiltration (29). These observations suggest that the C1/MES tumor types are inflammatory but protected from immune infiltration, suggesting an immune excluded tumor phenotype. Examining this subtype using the OV90 cell line with shRNA revealed UGDH as essential for spheroid viability, TIC viability and importantly, altered the TME in cocultures *in vitro* and in xenografts. Analysis of the gene expression from co-culture of OV90 knockdown in spheroids with mesothelial cells showed decreased expression of ECM components VCAN, LAMA3 and MMP1 and increased expression of differentiation and fibrosis markers CDH1 and FN1. Our findings align with previous reports of the effects of UGDH knockdown in cancer. In glioblastoma cell lines, silencing of UGDH with siRNA reduced viability and migration of cancer cells *in vitro* and tumor growth *in vivo*, largely due to the reduction of ECM proteins tenascin and laminin that promote glioblastoma progression (21). In breast cancer models, UGDH knockdown caused increased CDH1 and FN1 expression (26). The C1/MES tumor phenotype was replicated in the OV90 xenografts, with activated inflamed stroma observed in the OV90 negative control xenografts. Additionally, in line with what was observed in patients with the C1/MES subtype, overall survival improved in mice with UGDH knockdown OV90 xenografts compared to controls. An interesting phenotype of the OV90 knockdown tumors was the significantly impaired tumor growth compared to controls. Future studies will be done to investigate whether UGDH knockdown prevented tumors from establishing or if tumors were growing but regressed.

In contrast to the C1/MES subtype, the C4/DIF molecular subtype was described as having a low stromal response histologically and genetically, moderate immune infiltration in tumor and stroma and expression of markers of differentiation including E-cadherin, MUC16 and MUC1 (29,34). The low stromal activity in this subtype and low-moderate tumor immune infiltration suggests this tumor subtype is not inflamed or immune excluded and may represent a ‘cold tumor’. In this subtype, UGDH low expression was associated with a poorer prognosis. When we overexpressed UGDH in the C4/DIF ACI23 cell line, we observed increased spheroid formation but a reduced TIC population. Overexpression in ACI23 spheroids increased VCAN and LAMA3 expression, in opposition to what was observed with UGDH knockdown in OV90 spheroids. However, UGDH overexpression also increased TGF-B and CDH1 expression in ACI23 spheroids, and in co-culture with mesothelial cells. Like the knockdowns in OV90, there was reduced MMP1 expression compared to controls. Overexpression also increased cytokines IL-6, IL-8 and MCP-1 compared to controls. In the ACI23 xenografts we observed no significant difference in tumor size or necrosis, but interestingly, there was increased fibrosis in the UGDH overexpressed tumors compared to controls. These data suggest that UGDH overexpression in the C4 subtype activates the stroma, becoming more fibrotic. We did not have a syngeneic model of the C1/MES and C4/DIF subtypes to examine immune infiltration in xenografts, but our findings warrant further investigation to explore whether UGDH influences immune infiltration in EOC as was recently described in glioblastoma (51).

## Conclusions

UGDH expression in EOC influences the TME and reveals a distinct role for EOC-expressed UGDH in the C1/mesenchymal and C4/differentiated molecular subtypes of EOC. UGDH is a strong prospective therapeutic target in TICs, for the prevention or treatment of recurrent EOC especially in the mesenchymal subtype.

## Materials and Methods

### Antibodies and Reagents

Carboplatin (Cat# 2626) was purchased from Tocris Bioscience (Minneapolis, MN) and dissolved in ultra-pure water. Propidium Iodide (R37169) was from Thermo Fisher Scientific (Waltham, MA) and AnnexinV-FITC (556420) was from BD Biosciences (San Jose, CA). UGDH (HPA036656) was from Atlas Antibodies (Stockholm, Sweden) and GAPDH (MAB374) was from Millipore Sigma (Burlington, MA). Doxycycline used for *in vitro* studies was from Millipore Sigma (D5207, Burlington, MA). Inducible shRNA for knockdown of human UGDH (SMARTvector Inducible Lentiviral shRNA) and human UGDH for over-expression (Precision LentiORF) were purchased from Horizon Discovery (Cambridge, United Kingdom).

### Immunohistochemistry and Quantification

A TMA containing duplicate cores from archival samples of 96 HGS cases was generated as previously described (41). IHC staining for UGDH was performed using Novolink Polymer Detection Systems kit (RE7150-CE, Leica Microsystems, Mt Waverley, Australia) according to the manufacturer’s instructions. Briefly, slides were deparaffinized in xylene followed by graded alcohols then blocking for endogenous peroxidases and non-specific proteins (5 minutes at room temperature). Antigen retrieval was performed using Citrate Buffer pH 6.0 (005000, Thermo Fisher Scientific) at 110 °C for 15 minutes, followed by overnight incubation at 4 °C with the primary antibody (UGDH, 1:750). The secondary antibody and detection steps were performed using the Novolink Polymer Detection Systems Kit. Staining was scored by a pathologist (R.L) for intensity of staining and percentage of tumor cells expressing UGDH, providing an overall score of negative (score 0), weak (score 1), moderate (score 2) or strong (score 3). Four TMAs containing duplicate cores from 1: clear cell ovarian cancer, 2: mucinous ovarian cancer, 3: endometrioid ovarian cancer, 4: molecular subtyped ovarian cancer (Australian Ovarian Cancer Study, http://www.aocstudy.org/) were evaluated for expression of UGDH. IHC staining was performed at the Molecular Histopathology Laboratory (NCI, Frederick MD) on Leica Biosystems’ BondRX autostainer with the following conditions: Epitope Retrieval 1 (Citrate buffer) 20 min, UGDH (1:750, 30 min), and the Bond Polymer Refine Detection Kit (with omission of the Post Primary Reagent), (DS9800 Leica Biosystems Deer Park, IL,). Rabbit polyclonal isotype control (ab37415, Abcam Waltham, MA) was used in place of UGDH for the negative control. Slides were removed from the autostainer, dehydrated through ethanols, cleared with xylenes, and coverslipped. Positive control tissue included ovarian, prostate, and breast carcinoma tissue. Negative controls were performed for each TMA evaluated; negative controls include replacing the anti-UGDH antibody with nonspecific antibody of the same isotype (isotype control) taken from the same host. Slides were digitized with an Aperio ScanScope XT (Leica Microsystems, Buffalo Grove, IL) at 400X in a single z-plane. Aperio whole-slide images were evaluated and a threshold for positivity was determined using known positive controls by a board-certified pathologist. Cell detection algorithms were run to assess the positive cells for two separate outputs: cytoplasmic or membranous positive and nuclear positivity. Machine learning, random forest algorithms were trained for each tissue array to classify each cell detection as either epithelial or stromal; UGDH staining was separately quantified based on epithelial (tumor) or stromal. Stromal staining of UGDH was not observed, therefore only the epithelial/tumor staining expression was quantified. The staining intensity was scored using a scale of 0-3: 0 for no staining, 1 for mild staining, 2 for moderate, and 3 for strong staining and tumor H-score (42) was calculated using QuPath (52) as follows: H-score = (1 × (% cells 1+) + 2 × (% cells 2+) + 3 × (% cells 3+)).

### Cell lines and culture conditions

Ovarian cancer lines were obtained as gifts, or from ATCC or NCI-60 as described and were cultured as described (53). TIC-enriching spheroid culture conditions are previously described (30,47,50). Briefly, spheroids were generated by maintaining cells in ultra-low attachment (ULA) plates or flasks (Corning, Corning, NY) in defined medium. Experiments involving the TIC-enriched spheroid populations were grown for 3 days in defined medium in ULA plates before treatments were performed. LP3 mesothelial cells were obtained from the Coriell Institute and were grown in 1:1 Ham’s F12: Medium 199 containing 15% (v/v) FCS, penicillin (100 units per ml) and streptomycin (100 units per ml), 10ng/ml EGF and 0.4μg/ml hydrocortisone (Millipore Sigma, Burlington, MA). All cultures were maintained at 37°C in 5% CO_2_.

### Whole genome siRNA screen

The whole genome RNAi screen was performed at the Functional Genomics Lab (Rockville, MD), previously known as the Trans-NIH RNAi Facility (TNRF) as previously described (54,55). Briefly, the RNAi screen targeting 10,415 druggable genes (three individual siRNAs per gene) was conducted using OV90 cells and the Silencer^®^ Select Human Druggable Genome siRNA Library Version 4 (Ambion Thermo Fisher Scientific, Waltham, MA), in absence or presence of bardoxolone methyl. Adherent cells screening was carried out in 384-well white, solid, flat-bottom tissue culture plates (Corning, Corning, NY) while for spheroids screening 384-well black, clear, round-bottom ultra-low-attachment spheroid microplates were used (Corning, Corning, NY). Microplates were pre-stamped with one siRNA per well (2 μL, 400 nM) and, then 20ul of serum-free media containing Lipofectamine RNAiMax (Thermo Fisher Scientific, Waltham, MA) was added to each well. After 45 min incubation at room temperature, cells were added to wells in 20 μL media containing 20% FBS. Cells were cultured for 96 h, then cell viability was measured by the CellTiter-Glo Luminescent Cell Viability Assay (Promega, Madison, WI) with using EnVision Plate Reader (PerkinElmer, Boston, MA). Data analysis was performed as described (56). To rank genes that inhibited spheroid viability, the *Z*-score was calculated for each gene as: *Z=* (*x* - *μ)/σ, x* is the experimental value; *μ* is the median screen value; and *σ* is the standard deviation for the screen (57).

### RNA-sequencing alignment and analysis of ovarian cancer cell lines for molecular subtypes

Ovarian cancer cell lines were cultured in adherent conditions, and RNA was harvested according to the manufacturer’s instructions (74104, Qiagen, Germantown, MD). Sequencing was performed at the CCR Sequencing Facility (Leidos Biomedical Research, Frederick, MD). RNA-seq libraries were generated using TruSeq RNA Stranded Total RNA Library Prep Kits (TruSeq Illumina RS-122-2201) and sequenced on a total of 10 Hiseq 2500 lanes using the 125bp paired-end sequencing method (Illumina, San Diego, CA). Both reads of each sample were trimmed for adapters and low-quality bases using Trimmomatic software and aligned with reference human hg19 genome and ensemble v70 transcripts using Tophat software as stranded libraries. The sequencing quality of the reads was assessed per sample using FastQC (version 0.11.5) (http://www.bioinformatics.babraham.ac.uk/projects/fastqc/), Preseq (version 2.0.3) (58), Picard tools (version 1.119) (https://broadinstitute.github.io/picard/) and RSeQC (version 2.6.4) (http://rseqc.sourceforge.net/) (59). Reads were then trimmed using Cutadapt (version 1.14) (https://cutadapt.readthedocs.io/en/stable/) (60) prior to mapping to the hg19 human genome using STAR (version 2.5.2b) (https://github.com/alexdobin/STAR) (61) in two-pass mode. Overall expression levels were quantified using RSEM (version 1.3.0) (https://deweylab.github.io/RSEM/) (62). For normalization limma voom (version 3.48.3) (63) was used. For gene set enrichment, GSVA (64) was used using default parameters against 4 signatures from 4 subclusters (29) and used to create hierarchal clustering heatmap.

### Western blot analysis

Whole cell lysates were collected in lysis buffer: RIPA buffer (Thermo Scientific, Waltham, MA) containing 1× protease inhibitor (78430, Thermo Fisher Scientific, Waltham, MA) 1× phosphatase inhibitor (Phos-STOP, PHOSS-RO, Millipore Sigma, Burlington, MA). After a brief incubation on ice, the lysates were homogenized by passing the samples through 26-G needles, followed by centrifugation at 16,000 g, 4°C, for 20 min to collect the supernatant. Protein concentration was quantified by microbicinchoninic acid assay (23227, Thermo Fisher Scientific, Waltham, MA). Lysates (30 μg) were separated by SDS-PAGE under reducing conditions, transferred to nitrocellulose membranes, and blocked in Intercept (TBS) blocking buffer (927-66003, LI-COR Biosciences, Lincoln, NE). Membranes were incubated with primary antibodies diluted in blocking buffer overnight at 4°C, (UGDH 1:1000), (GAPDH 1:10000), washed with Tris-buffered saline containing 0.1% Tween 20 (TBST), then incubated with fluorescent secondary mouse or rabbit IgG antibodies (IRDye, LI-COR Biosciences, Lincoln, NE). Images were generated using the Odyssey system and software (LI-COR Biosciences, Lincoln, NE).

### Cell viability

Cell viability was assessed as previously described (30,50) using CellTiter-Glo (Promega, Madison, WI) according to manufacturer’s instructions.

### Sphere formation

Sphere formation was performed as previously described (30,50). OV90 cells were seeded at 2000 cells/well in 96-well ULA plates (3474, Corning, NY), in TIC-enriching medium (TEM) with 1μg/mL doxycycline for 7 days, fresh culture medium containing growth factors was replenished every 48 hours. ACI23 cells were seeded at 1000 cells/well in 96-well ULA plates in TEM for 7 days, fresh culture medium containing growth factors was replenished every 48 hours. After 7 days the spheroids were incubated with DRAQ5 (62254, Thermo Fisher Scientific, Waltham, MA, USA) at 1μM for 15 minutes prior to imaging as described (50). Quantification of spheroids was performed using NIS Elements software (Nikon, Melville, NY), as described (50) and the number of spheroids measuring an area of >1000 μm^2^ were counted.

### Colony formation

The colony formation assay was performed according to the manufacturer’s instructions (CBA-130, Cell Biolabs, San Diego, CA). Briefly, a base layer of agar was plated and allowed to solidify, before adding a cell-agar layer. The agar layers were topped up with appropriate media, doxycycline-containing media for experiments involving inducible shRNA. Culture media was refreshed every 72 hours, and following the lysis protocol, fluorescence was measured using a plate reader.

### Cytokine Array

Cytokine analysis was performed on cell culture supernatants using LEGENDplex™ HU Essential Immune Response Panel (740930, Biolegend, San Diego, CA) according to the manufacturer’s instructions (65).

### Flow cytometry

ALDH enzymatic activity was assessed as previously described (30,47,50), using ALDEFLUOR (Stem Cell Technologies, Seattle, WA) according to the manufacturer’s instructions. Following ALDH staining, cells were incubated with CD133-APC antibody (BD Biosciences, Ashland, OR) at 1:20 dilution in ALDEFLUOR buffer for 25 minutes on ice, protected from light. Cells were washed in PBS and resuspended in 400μl PBS for analysis on a BD FACSVerse cell analyzer (BD Biosciences, Franklin Lakes, NJ). Cell death was assessed by Annexin V (640905, Biolegend, San Diego, CA) and propidium iodide (PI) (R37169, Thermo Scientific, Waltham, MA) staining on cells treated as indicated, as previously described (30,50).

### Quantitative real-time PCR (qRT-PCR)

Total RNA of mesothelial cells from co-culture was extracted using RNeasy Mini Kit according to the manufacturer’s instructions (74106, Qiagen, Mansfield, MA). Total RNA was extracted from frozen xenograft tumors using TRI Reagent (AM9738, Thermo Scientific, Waltham, MA) according to the manufacturer’s instructions, immediately following overnight thawing in RNAlater™-ICE Frozen Tissue Transition Solution (AM7030, Thermo Scientific, Waltham, MA). RNA was converted to cDNA using High-Capacity cDNA Reverse Transcription Kit according to the manufacturer’s instructions (4368814, Applied Biosystems, Thermo Scientific, Waltham, MA). TaqMan™ Array Human Extracellular Matrix & Adhesion Molecules (4414133, Applied Biosystems, Thermo Scientific, Waltham, MA) and TaqMan™ Gene Expression Assays for hMMP1 (Hs00899658_m1), hFN1 (Hs01549976_m1), hLAMA3 (Hs00165042_m1), hVCAN (Hs00171642_m1), hTGFB1 (Hs00171257_m1), hTIMP3 (Hs00165949_m1), hTNC (Hs01115665_m1), hCOL1A1 (Hs00164004_m1), hCDH1(Hs01023895_m1), hIL6 (Hs00174131_m1), hIL-8 (CXCL8) (Hs00174103_m1), hCCL2 (Hs00234140_m1), were used with TaqMan™ Fast Advanced Master Mix (4444963, Applied Biosystems, Thermo Scientific, Waltham, MA) and qRT-PCR was performed using ViiA 7 System. The comparative threshold cycle (Ct) method was used to calculate the relative gene expression and target genes values were normalized to the expression of the endogenous reference gene.

### *In vivo* studies

All animal studies were approved by the NCI Animal Care and Use Committee, IACUC Number MOB-025-1. Intra-bursal xenografts were generated by injection of 0.5 x10^5^ cells in 5 μL PBS into the right ovarian bursa of 8-week-old female athymic Nu/Nu mice. For controls, 5 μL PBS was injected into the left ovarian bursa of each mouse. For tumor burden studies, both ovarian bursa were injected with 2.5×10^5^ cells. Mice injected with OV90 cells containing the DOX-inducible shRNA were fed doxycycline chow (200mg/kg, S3888, Bio-Serv, Flemington, NJ) for the duration of the study. The animals were monitored for health and survival in days was recorded as mice met NIH Animal Care and Use Committee-approved humane criteria for euthanasia.

### Statistical Analysis

*In vitro* assays were performed in triplicate on three independent occasions and were analyzed with t-tests or one-way ANOVA with post-tests where applicable. Results are presented as mean ± SEM with p values ≤ 0.05 considered significant. Kaplan-Meier analysis was used to analyze overall survival and progression-free survival for IHC analyses, and Mantel-Cox log-rank was used to compare groups. Statistical analyses were performed using Prism 8.0 software (GraphPad, San Diego, CA, USA).

## Author Contributions

The study was conceptualized by BSH, CMA. Investigation and formal analysis were performed by BSH, RK, MWC, SK, FN, CDH, LFH. Immunohistochemistry and resources by DB, EE, MG, RR, RL, NT, JDH, DDLB. Software by JS, BT, FE, AA, MLN, KMM. Original draft was written by BSH, all authors read and edited the final draft. All authors have read and agreed to the published version of the manuscript.

## Acknowledgments

The authors are grateful for the technical assistance from the National Cancer Institute microscopy core facility staff. Additionally, we thank Elena Kuznetsova and Geneti Gaga for their expertise with intrabursal injections.

